# Predator-Induced Tail Coloration Toward Diverse Dragonfly Nymphs in Tadpoles of the East Japan Tree Frog (*Dryophytes leopardus)*

**DOI:** 10.1101/2025.01.13.632661

**Authors:** Akihiro Noda, Katsutoshi Watanabe

**Affiliations:** Department of Zoology, Graduate School of Science, Kyoto University, Kitashirakawa-oiwake-cho, Sakyo-ku, Kyoto 606-8502, Japan

## Abstract

Anuran larvae can change their coloration and morphology to avoid predation. Particularly, some tadpoles in the genus *Dryophytes* in the Americas develop deep tail fins and bright orange tail coloration in response to predators such as dragonfly nymphs. These conspicuous tails are hypothesized to attract predator attacks, protecting vital body parts from fatal injuries. However, it remains unclear whether the extent of the induced phenotypic changes differs between various predators with distinct feeding patterns and habitats. To explore this, we reared tadpoles of the East Japan tree frog, *Dryophytes leopardus*, with four dragonfly species, water bug, water scorpion, or newt, and measured phenotypic changes, especially focusing on tail coloration. We found that the aeshnid *Anax nigrofasciatus*, a sit-and-wait predator that climbs on aquatic vegetation, induced bright orange and broad tails of tadpoles. Other odonate species that forage in and on the substrate also elicited tail coloration or deeper larval tails, but it took more time to induce the tadpoles’ response compared to *A. nigrofasciatus*. In contrast, non-odonate predators did not induce color and morphological changes. This study provides the first evidence of predator-induced tail coloration in an Asian species of *Dryophytes*, which appears to be a defensive response specifically to dragonfly nymphs, as visually adept predators. Moreover, the differences in the extent and timing of the phenotypic changes among odonate species suggest that anti-predator phenotypes are modulated based on relative predation risks, considering the predators’ foraging patterns and visual capabilities, thereby allowing tadpoles to minimize physiological costs and unintended risks associated with the induced traits.

---

The utilization of prey’s body color and shape is highly dependent on its context, including the sensory capabilities of predators and the background condition (e.g., Espanha et al., 2016; Barnett et al., 2018). Especially in aquatic habitats, substantial variation exists in background conditions and the composition and abundance of predators across different localities, making it difficult for prey to achieve a uniform predation avoidance ability across their range (Van Buskirk, 2002; McIntyre et al., 2004). Therefore, such environmental uncertainty has generally been believed to favor plastic rather than constitutive anti-predator defenses (Agrawal and Karban, 1999; McIntyre et al., 2004).

Anuran larvae, or tadpoles, have served for decades as a model system for investigating induced defense mechanisms, as well as predator avoidance strategies and trade-offs mediated by plastic traits (Kasumovic, 2013; Kruger and Morin, 2020). They often rely on chemical cues to evaluate predation risk through continually released predator-borne cues and digestion-released, prey-borne cues, and then exhibit predator-induced defense mechanisms (Brönmark and Hansson, 2000; Hettyey et al., 2010, 2015). Predator-induced plastic responses in tadpoles can be categorized into four primary domains: behavioral, morphological, physiological, and chemical defenses. Behavioral responses encompass changes in overall activity levels (e.g., Lawler, 1989; Takahara et al., 2012a; Reuben and Touchon, 2021), increased burst swimming speed (Takahara et al., 2008), and spatial avoidance of predators (e.g., Laurila et al., 1998; Nunes et al., 2013). Morphological changes for predator avoidance have been well documented in tadpoles of various species (Relyea, 2001; Van Buskirk, 2002; Richter-Boix et al. 2007). For instance, tadpoles undergo changes in body size (i.e., correlated with growth rate and metamorphosis timing), body shape such as the development of deeper tail fins (Lardner, 1998, 2000), and coloration (McCollum and Van Buskirk, 1996; Touchon and Warkentin, 2008). Behavioral and morphological responses are sometimes co-expressed (Hossie et al., 2017). Physiological responses include changes in respiration rate and gut evacuation rate (Steiner, 2007; Steiner and Van Buskirk, 2009; Costello and Michel, 2013). These may arise due to trade-offs in predator avoidance (Relyea and Auld, 2004) or as responses to partially offset the costs of lower energy intake resulting from evasion (Barry and Syal, 2013). Additionally, tadpoles of certain groups, particularly toads, evade predation through toxin production (Bókony et al., 2016; Hettyey et al., 2019).

The types and intensities of induced responses in tadpoles often differ according to predator types (Relyea, 2001; Teplitsky et al., 2004, 2005; Hettyey et al., 2011). For instance, Kishida and Nishimura (2005) revealed that tadpoles of *Rana pirica* developed deeper tail fins in response to dragonfly nymphs, whereas in the presence of salamander larvae, they exhibited not only deeper tails but also bulging heads and bodies. Tadpoles expressing these predator-specific traits should have a survival advantage over those with mismatched or non-induced phenotypes when faced with predation. Conversely, expressing such defense traits incurs energetic costs and can increase vulnerability to non-focal predators (Touchon and Warkentin, 2008; Innes-Gold et al., 2019). Due to these costs, defense traits are expected to remain plastic rather than fixed, and their types and intensities can vary depending on predator species, sometimes exhibiting predator-specific responses.

Dragonfly nymphs, as one of the major predators of tadpoles, are known to induce morphological responses characterized by deep tail fins across many anuran species (Relyea and Werner, 2000; Van Buskirk, 2002). This induced broad tail has been repeatedly hypothesized to improve the swimming performance of tadpoles and facilitate their escape from predators (Smith and Van Buskirk, 1995; McCollum and Leimberger, 1997; Van Buskirk and McCollum, 2000a). However, experimental evidence of this benefit is lacking (Van Buskirk et al., 2003; Johnson et al., 2008). Moreover, a few tree frog species in the genus *Dryophytes* respond to odonate predators by not only exhibiting deeper tails but also turning their entire tails bright orange (McCollum and Van Buskirk, 1996; Van Buskirk and McCollum, 1999; LaFiandra and Babbitt, 2004). Thus, as another function, the induced deep and brightly colored fin is thought to serve as a lure, diverting predator attacks away from the more vulnerable head and body regions (Van Buskirk et al., 2003, 2004; Hawlena et al., 2006; Johnson et al., 2008). Given that tail fins tear easily (Doherty et al., 1998) and that tadpoles can maintain their swimming performance even after partial tail loss (Hoff and Wassersug, 2000; Van Buskirk and McCollim, 2000b), the tail is considered expendable relative to other body parts. Because dragonfly nymphs are predominantly visual predators, exhibiting the ability to discern both the shape and color of immobile as well as moving prey (Bybee et al., 2012; Espanha et al., 2016), color and shape changes in tadpoles are likely to be effective in avoiding predation by odonates.

Among dragonfly nymphs, several ecological types have been identified based on microhabitat usage and foraging behavior (Pritchard, 1965; Corbet, 1999), including climbers (e.g., Green darner, *Anax junius*), moving among aquatic vegetation and ambushing prey on plants; sprawlers (e.g., Globe skimmer, *Pantala flavescens*; Sherratt and Harvey, 1989), inhabiting pond bottoms and swimming through the water column to attack; and burrowers (e.g., Golden-ringed dragonfly, *Cordulegaster boltonii*), burrowing in mud or bottom debris and often lying in wait for prey with only the eyes protruding. The reliance on vision for foraging varies among these types (Pritchard, 1966; Corbet, 1999). For tadpoles, the frequency of encounters with and relative predation risk of these predators differ among habitats and seasons. These differential predation risks likely determine the cost–benefit balance of induced traits, thereby suggesting that tadpoles may exhibit different defense responses. However, prior experimental studies have primarily focused on a single dragonfly species and have not examined how tadpole responses may vary among odonate (and other) species.

The East Japan tree frog, *Dryophytes leopardus*, which was recently recognized as distinct from *Dryophytes japonicus*, is a prevalent species with a wide distribution encompassing northeastern Japan (central to eastern Honshu and Hokkaido), as well as regions outside Japan including Sakhalin Island (Shimada et al., 2025). Tadpoles of this species inhabit various environments such as rice paddies, marshes, ponds, puddles, and other ephemeral water bodies (Matsui and Maeda, 2018). Tadpoles, including this species, serve as important prey for numerous aquatic predators, including dragonfly nymphs, water scorpions, and red-bellied newts (Ohba and Nakatsuji, 2006; Hirai et al., 2020; Kita et al., 2020; Ozaki and Nishikawa, 2023). Morphological responses to predators have also been documented in this species, with dragonfly nymphs inducing deep tail fins, while chemical cues from goldfish prompt the development of shallow fins (Takahara et al., 2012b). The occurrence of orange tail coloration has also been briefly reported, although it has been attributed to ingested algae rather than predator-induced plasticity (Doi et al., 2005; Yuasa, 2006, 2007). Despite being rarely documented, tadpoles exhibiting brightly colored tails were frequently observed at our study site in Kyoto, central Japan (Noda, personal observation).

To clarify the cause and function of the orange tail coloration in tadpoles of *D. leopardus*, we conducted laboratory experiments with the tadpoles and several predator species. These experiments aimed to achieve two primary objectives. First, we investigated whether the tail coloration of tadpoles is induced by dragonfly nymphs, and if so, whether the intensity and speed of this response vary among different odonate species with different ecological characteristics. Simultaneously, we examined changes in body shape that are known to be plastically induced by predators. The second objective was to assess whether tadpoles exhibit color/morphological responses to predators characterized by different foraging styles compared to odonates. We used water bugs (*Appasus major*), water scorpions (*Laccotrephes japonensis*), and newts (*Cynops pyrrhogaster*) as non-odonate predators. Water bugs and scorpions capture prey by enclosing them with their frontal legs, and newts swallow prey whole. Accordingly, predation avoidance strategies involving tail coloration, which are based on diverting attacks toward a specific body part, are expected to be less effective against these predators than against dragonfly nymphs. Therefore, tadpoles may exhibit weakened or no color/morphological responses to minimize the costs associated with developing plastic traits. In both experiments, tadpoles were reared with a single predator species, and their morphological traits of interest were compared with those of tadpoles raised in predator-free tanks. By analyzing the extent of tail coloration in tadpoles exposed to diverse predators, we revealed that tadpoles modulate their induced responses according to the risks posed by predators with different foraging types and sensory capabilities.

## MATERIALS AND METHODS

Two series of laboratory experiments were carried out: Experiment I involved the comparison of induction of color/morphological changes in tadpoles by four different dragonfly species as predators, and Experiment II employed three non-odonate predators.

### Experiment I

Seven mating pairs of *Dryophytes leopardus* were captured in the paddy fields at Kyoto University (35°01’N, 135°47’E) on June 9 and 10, 2022. Following capture, mating pairs were housed overnight in plastic cages (19 × 19 × 14 cm; 6 cm water depth), and eggs were obtained the following day. The eggs hatched in about 2–3 days, and after hatching, tadpoles were separately kept in plastic containers (43 × 31 × 16 cm; 6 cm water depth), as those from the same clutch were kept together in a single container. A room temperature of 20°C was maintained until the experiments began, and tadpoles were fed powdered food (Kyorin Co., Ltd.) every two days. In this experiment, four species of dragonfly nymphs were used as predators taking into account their foraging styles: *Anax nigrofasciatus* (Black-striped lesser emperor; climber), *Gynacantha japonica* (Blue-spotted dusk-hawker; climber), *Orthetrum albistylum speciosum* (White-tailed skimmer; burrower), and *Pantala flavescens* (Wandering glider; sprawler). Nymphs of *A. nigrofasciatus* collected in Sakura City, Chiba Prefecture, and those of *G. japonica* collected in Higashihiroshima City, Hiroshima Prefecture, were obtained from an online vendor, and *O. a. speciosum* and *P. flavescens* were collected from the same paddy field as the mating frog pairs. The dragonfly nymphs were maintained individually in plastic trays (23 × 17 × 8 cm) and fed tadpoles of *D. leopardus* every two days until the beginning of the induction experiment.

The induction experiment commenced on July 12, 2022. Each experimental tank housed 21 tadpoles, with three individuals randomly selected from each of the seven egg clutches to randomize genetic effects in each tank. Tadpoles in a tank were assigned to one of five treatments (Ax group with *A. nigrofasciatus*; Gy group with *G. japonica*; Or group with *O. a. speciosum*; Pf group with *P. flavescens*; and the control Ct1 group) with eight replicates per treatment. Each experimental tank (31 × 18 × 24 cm; 10 cm water depth) was equipped with a transparent isolation case (17 × 9 × 10 cm) with approximately 5 mm diameter holes spaced about 5 mm apart, where one dragonfly nymph was introduced. This setup allowed visual and chemical-mediated interaction between the predator and tadpoles without physical contact. The tanks were aerated and maintained under a 14-h light/10-h dark photoperiod at a room temperature of 22°C, with half of the water in all tanks changed every three days. Air was continuously supplied to the isolation case within each tank through an aquarium air stone. Tanks designated for the Ct1 group lacked predators and utilized an empty isolation case. Tadpoles in the experimental tanks were fed every two days with commercial food for Axolotl, *Ambystoma mexicanum* (Kyorin Co., Ltd.), while dragonfly nymphs received two tadpoles as food every two days.

### Experiment II

The design of Experiment II mirrored that of Experiment I, with adjustments made for sampling dates, the interval between hatching and the experiment’s commencement, and the selection of predators. For this experiment, three mating pairs of *D. leopardus* were captured from the paddy fields at Kyoto University on May 18 and 20, 2022. Eggs were collected and tadpoles from clutches were reared in the same way as in Experiment I until the experiments began. Three predator species were chosen based on their foraging style and prey capture behavior: Ferocious water bug, *Appasus major* (sit-and-wait predator that seizes prey with its frontal legs), water scorpion, *Laccotrephes japonensis* (sit-and-wait predator that grasps prey with its frontal legs), and Japanese fire belly newt, *Cynops pyrrhogaste*r (active tracker that swallows prey whole). Aquatic insects, collected in Higashihiroshima City, were purchased from the online vendor, while red-bellied newts were collected from irrigation ditches in Kyoto City (35°05’N 135°47’E). These predators were individually housed in plastic trays and fed with tadpoles every two days.

The induction experiment began on June 18, 2022. Each experimental tank housed 21 tadpoles, with seven individuals randomly selected from each of the three egg clutches. Tadpoles in each tank were allocated to one of four treatments (Be group with *A. major*; Ws group with *L. japonensis*; Fn group with *C. pyrrhogaster*; and the control Ct2 group) with eight replicates per treatment. One predator was introduced into an isolation case in each tank. Other experimental setups, i.e., tank design, room temperature, light condition, and feeding frequency, were kept the same as in Experiment I.

### Measuring Coloration and Morphological Responses

All tadpoles were photographed two and three weeks after the start of the experiment using a Canon🄬 EOS RP mirrorless camera. Individual tadpoles were placed in a narrow acrylic aquarium (20.5 × 4.5 × 13.5 cm) with a ruler and color chart in the background. To minimize the effect of posture variation among individuals, multiple photographs were taken per tadpole, and the image most suitable for measurement was selected. Each trait was measured once per individual. The distance between the aquarium and the camera was kept constant for all individuals. All photos were taken with a F/16 aperture, a 1/20th second exposure time, and ISO 3200. Consistency in artificial lighting conditions was maintained for individuals photographed on the same day, with the white balance manually set and standardized at 4600 K. Some individuals were excluded from the analysis due to poor nutritional condition or significant tail loss during rearing. The exclusion criteria included: (1) head and body length less than 6 mm, (2) severe tail damage or loss, and (3) inability to maintain posture in water.

Tail fin parts, defined as all tail areas excluding the central tail muscle, were trimmed manually using the image measuring software, Natsumushi version 1.12.2 (Tanahashi and Fukatsu, 2018; https://sites.google.com/site/mtahashilucanid/program/natsumushi) and selected for color measurement. The color of the tail was quantified using the average of CIELAB values (CIE 1976) calculated in Adobe Photoshop 25.5.1 (Adobe Inc.). In the CIELAB color space, color was represented using three components: the lightness component L* and the color components a* and b*. The L* component ranged from black to white between 0–100, while a* and b* components ranged from green to red and blue to yellow, respectively, on a scale of −128 to 127. As Photoshop displays images in 8-bit format, these values were rescaled to range from 0 to 255. Therefore, L* was multiplied by 100/255 and a* and b* were adjusted by subtracting 128, respectively, to facilitate analysis (Fig. 1). Due to incorrect white balance, photographs taken at the second week of Experiment II were excluded from the analysis of tail coloration. During the third week of Experiment II, photographs of 11 individuals from the Ct2 group were also taken with an incorrect white balance, so the color tone was later corrected using Photoshop. We used the “Match Color” function in the “Image Adjustments” menu, referencing one of the other photographs from the Ct2 group that was taken under the same conditions, with the white balance properly set to 4600 K. Additionally, morphological measurements were conducted based on the photographs using Natsumushi. Morphological traits included tadpole body length (BL), tail length (TAL), and tail depth (TD) were measured using the scale within each image.

**Fig. 1.**
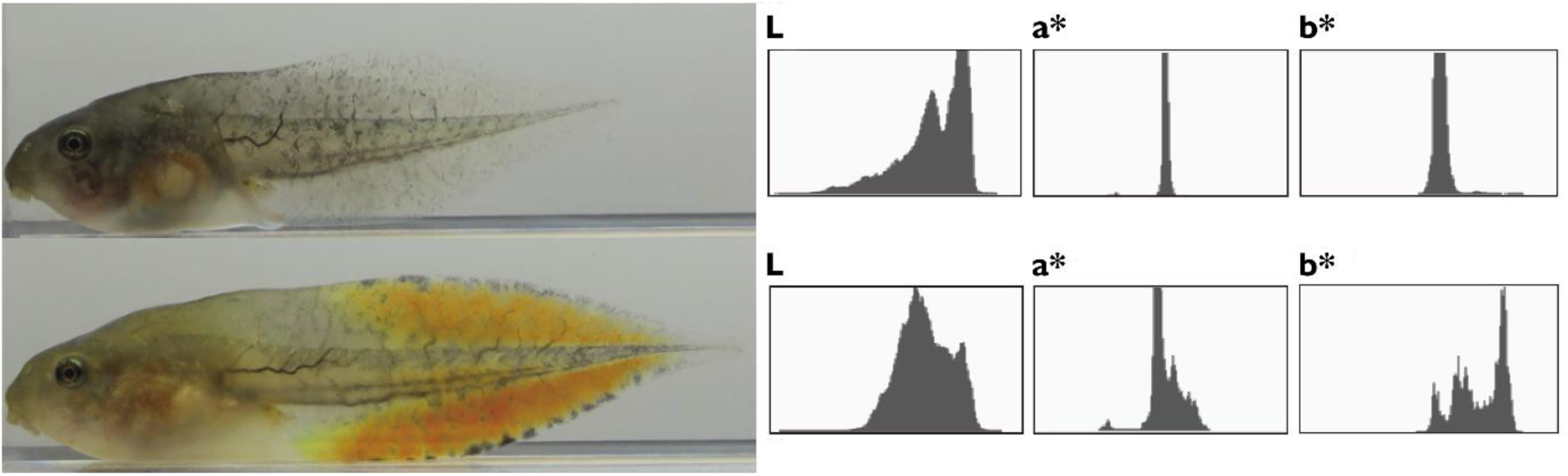
Photographs of tadpoles of *Dryophytes leopardus* reared without predator (top) and with *Anax nigrofasciatus* (bottom) and their corresponding Photoshop Lab color histograms. The mean values of tail coloration are L = 61.2, a* = -2.5, and b* = 7.96 (without predator), and L = 57.96, a* = 1.75, and b* = 35.51 (with *A. nigrofasciatus*), respectively.

### Statistical Analyses

All statistical analyses were conducted in R version 4.3.2 (R Core Team, 2023). Linear mixed models (LMMs) were employed based on the measured traits, using the lmerTest package (Kuznetsova et al., 2017) to estimate differences between the control group and the other treatments. The experimental tank was treated as a random factor in the LMMs. When color components (e.g., a* and b*) were utilized as the response variables, the treatment (predator) was used as the explanatory variable in the models to compare the treatment groups to the control group. In the same way, when body length (BL) was utilized as the response variables, the treatment was used as the explanatory variable in the models. On the other hand, in analyses for TAL and TD as the response variable, four models were constructed for each trait. These models included: the model with only BL as the explanatory variable, the model with only the treatment as the explanatory variable, the model with both BL and treatment as explanatory variables, and the model incorporating BL, treatment, and their interactions. The final model was determined via backward elimination, by applying likelihood ratio (LR) tests. The significance level was set at 5%. If a significant difference was found between the former and reduced models, the former model was adopted as the final model for further analysis.

After constructing final models, we evaluated the overall effect of predator treatment by performing ANOVA via the anova function in the lmerTest package. Subsequently, comparisons between each treatment group and the control were performed using Dunnett’s test with estimated marginal means calculated by the emmeans package (Lenth, 2025). Marginal means were derived under the asymptotic framework. The significance level was set at 5%, with treatment groups considered significantly different from the control group if the *P* values were below this threshold. Orange tail coloration and increased tail depth were reported as predator-induced traits in previous studies (McCollum and Van Buskirk, 1996; Touchon and Warkentin, 2008), so we decided to focus primarily on b* values and tail depth. To evaluate the covariation between tail coloration and morphology at the third week of the experiment, we also conducted Pearson’s correlation tests between residuals of b* values and tail depth (TD), after removing the effects of body length (BL) via linear regression. This analysis was performed separately for each predator treatment that showed a significant difference from the control group in either trait.

## RESULTS

### Coloration

Experiment I showed that predator presence induced changes in tadpole tail coloration. At the second week, the overall effects of predator treatment were significant for both a* and b* values (a*: *F* = 11.41, *P* < 0.001; b*: *F* = 12.05, *P* < 0.001). The response was strongest with *Anax nigrofasciatus* (Ax group), where tadpoles developed significantly more orange tails than controls, as indicated by a consistent decrease in a* and increase in b* values (Dunnett’s test, *P* < 0.05; Table 1; Fig. 2). At the third week, a significant overall effect of predator treatment remained only in b* values (*F* = 11.31, *P* < 0.001), not in a* values (*F* = 2.18, *P* = 0.096). Nevertheless, tadpoles in the Ax group still showed significantly lower a* values (*P* = 0.028) and higher b* values (*P* < 0.001) than controls. A few individuals in the Ax group exhibited particularly high values in both a* and b*, corresponding to intensely vivid orange tails. These data points likely represent the upper range of predator-induced color expression, rather than artifacts. A weaker, yet still significant, effect was observed with tadpoles exposed to *Orthetrum albistylum speciosum* (Or group), which showed elevated b* values compared to the control (*P* = 0.045). In the tadpoles with *Pantala flavescens* (Pf group), a similar trend was noted, but the differences from the control group were not statistically significant (*P* = 0.143 in b* value). In contrast to the results of Experiment I, overall treatment effects in Experiment II were not significant (a*: *F* = 0.49, *P* = 0.689; b*: *F* = 1.84, *P* = 0.166), and no group showed significant differences from the Ct2 group in the a* or b* values at the third week (Table 2; Fig. 3).

**Fig. 2.**
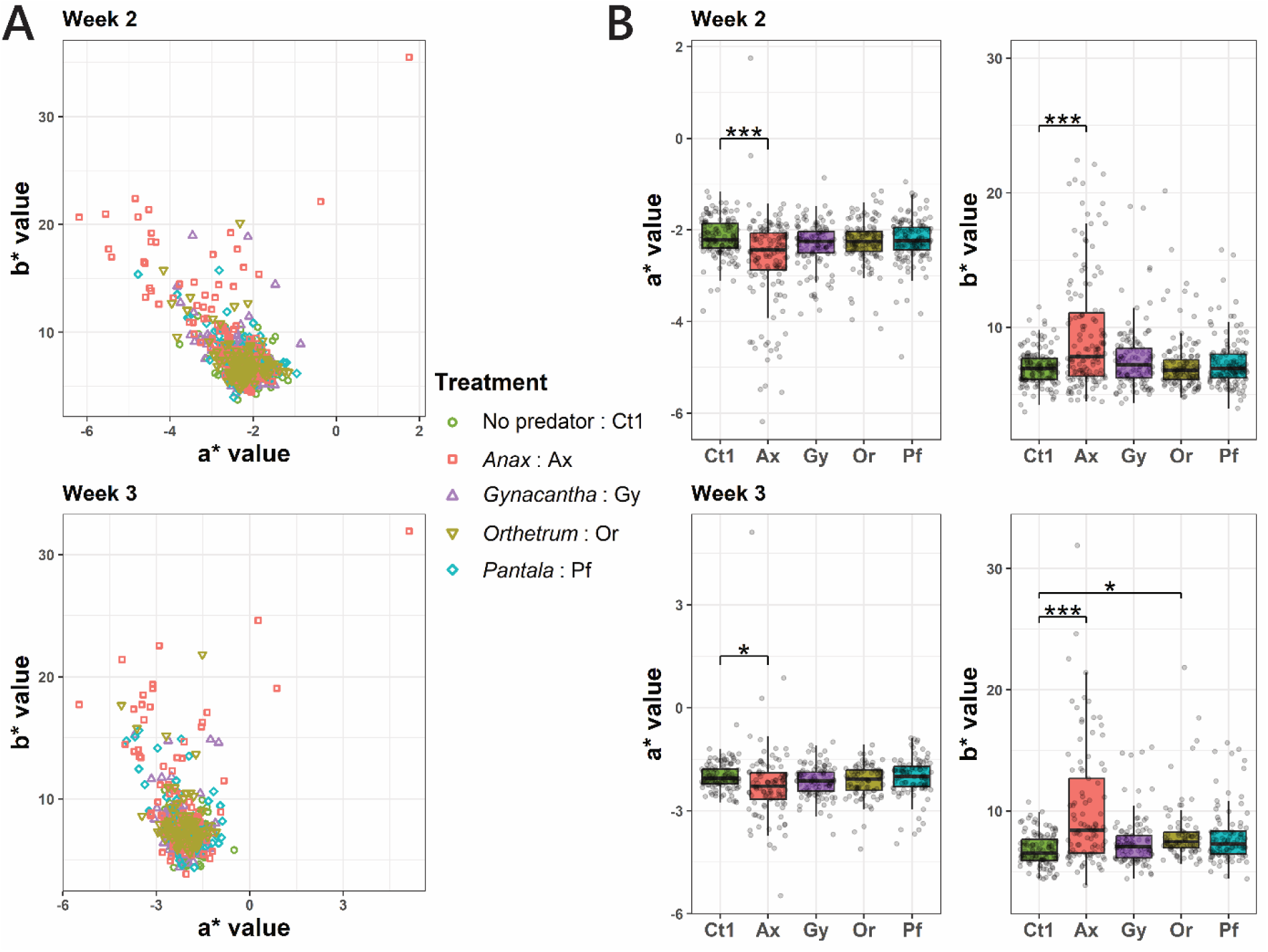
(A) Scatterplots of tail coloration parameters (a* and b* values) of tadpoles two weeks (top) and three weeks (bottom) after the start of Experiment I, using different dragonfly species. Tadpoles were reared in the presence of no predators (the control group; Ct1), *Anax nigrofasciatus* (Ax), *Gynacantha japonica* (Gy), *Orthetrum albistylum speciosum* (Or), or *Pantala flavescens* (Pf). (B) Box-and-whisker plots displaying the range of the tail color traits (a* and b* values) two weeks (top) and three weeks (bottom) after the start of Experiment I. The asterisks in the figure indicate that there were significant differences compared to the control group, which were led by Dunnett’s test (see Materials and Methods). \**P* < 0.05, \*\*\**P* <0.001.

**Fig. 3.**
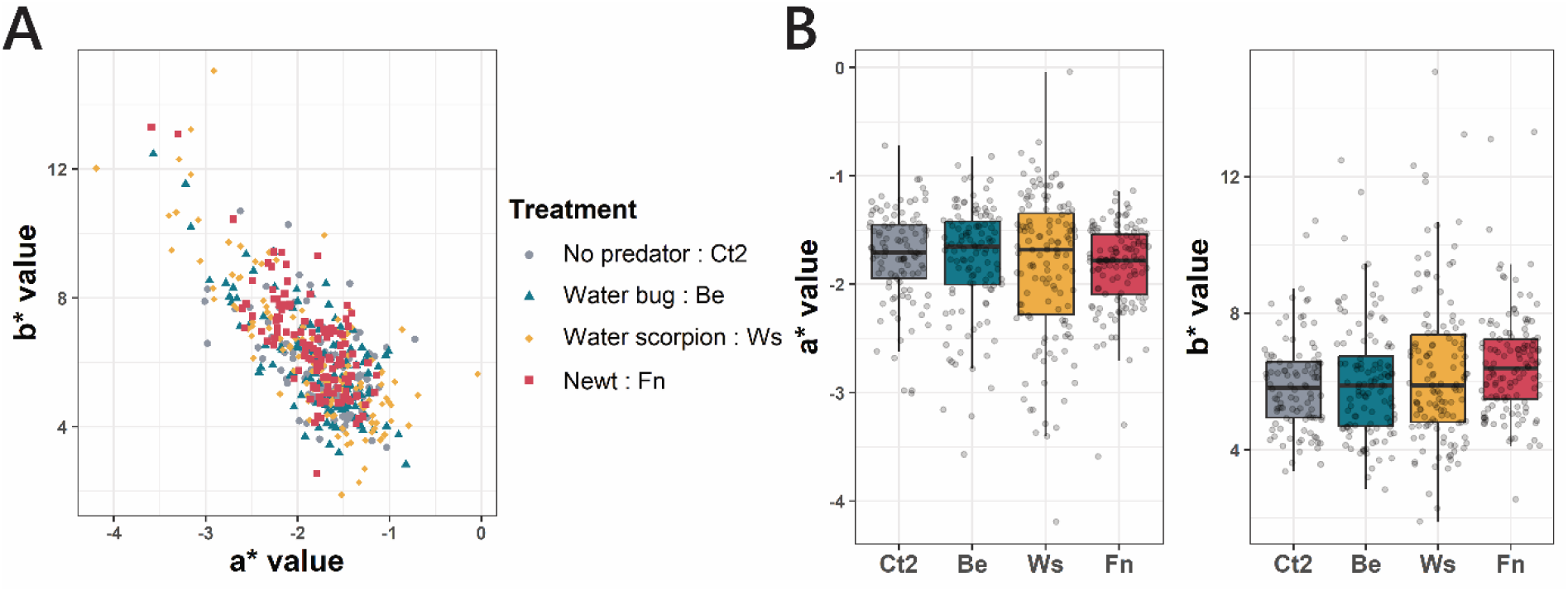
(A) Scatterplots of tail coloration parameters (a* and b* values) of tadpoles three weeks (bottom) after the start of Experiment II, using non-odonate species. Tadpoles were reared in the presence of no predators (the control group; Ct2), water bug (*Appasus major*; Be), water scorpion (*Laccotrephes japonensis*; Ws), and fire belly newt (*Cynops pyrrhogaster*; Fn). (B) Box-and-whisker plots displaying the range of the tail color traits (a* and b* values) three weeks after the start of Experiment II.

**Table 1.** Summary of model results for tail coloration (a* and b* values) measured at Week 2 and Week 3 after the start of Experiment I. Tadpoles were reared in the presence of no predators (the control group), *Anax nigrofasciatus* (Ax), *Gynacantha japonica* (Gy), *Orthetrum albistylum speciosum* (Or), or *Pantala flavescens* (Pf). Fixed effects include the corresponding estimates (Est.), standard errors (SE), degrees of freedom (df), t-values (*t*), and model P-values (*P*_m_). Post hoc comparisons were performed using Dunnett’s test to compare each predator treatment to the control group (*P*_D_). Variance (Var.) and standard deviation (SD) for the random effect (Tank) and residuals are also shown. The control group served as the reference level in all models.

Week 2
| Effect | Est. | SE | df | $t$ | $P_m$ | $P_D$ |
| --- | --- | --- | --- | --- | --- | --- |
| (Intercept) | -2.158 | 0.052 | 32.153 | -41.753 | <0.001 |  |
| **Ax** | -0.462 | 0.074 | 33.245 | -6.256 | <0.001 | <0.001 |
| **Gy** | -0.157 | 0.076 | 37.975 | -2.062 | 0.046 | 0.129 |
| **Or** | -0.112 | 0.076 | 37.689 | -1.462 | 0.152 | 0.392 |
| **Pf** | -0.074 | 0.075 | 34.351 | -0.991 | 0.329 | 0.689 |

| Random Effect | Var. | SD |
| --- | --- | --- |
| Tank | <0.001 | 0.009 |
| Residual | 0.378 | 0.615 |

$b^*$ value Week 2
| Effect | Est. | SE | df | $t$ | $P_m$ | $P_D$ |
| --- | --- | --- | --- | --- | --- | --- |
| (Intercept) | 7.048 | 0.296 | 31.896 | 23.786 | <0.001 |  |
| **Ax** | 2.568 | 0.422 | 32.707 | 6.083 | <0.001 | <0.001 |
| **Gy** | 0.620 | 0.431 | 35.535 | 1.439 | 0.159 | 0.405 |
| **Or** | 0.219 | 0.431 | 35.478 | 0.507 | 0.615 | 0.929 |
| **Pf** | 0.299 | 0.425 | 33.461 | 0.705 | 0.486 | 0.849 |

| Random Effect | Var. | SD |
| --- | --- | --- |
| Tank | 0.269 | 0.519 |
| Residual | 7.679 | 2.771 |

Week 3
| Effect | Est. | SE | df | $t$ | $P_m$ | $P_D$ |
| --- | --- | --- | --- | --- | --- | --- |
| (Intercept) | -1.998 | 0.063 | 25.632 | -31.725 | <0.001 |  |
| **Ax** | -0.247 | 0.093 | 26.952 | -2.659 | 0.013 | 0.028 |
| **Gy** | -0.141 | 0.091 | 27.399 | -1.558 | 0.131 | 0.338 |
| **Or** | -0.145 | 0.097 | 32.740 | -1.495 | 0.145 | 0.373 |
| **Pf** | -0.035 | 0.094 | 26.841 | -0.369 | 0.715 | 0.966 |

| Random Effect | Var. | SD |
| --- | --- | --- |
| Tank | 0.002 | 0.043 |
| Residual | 0.429 | 0.655 |

Week 3
| Effect | Est. | SE | df | $t$ | $P_m$ | $P_D$ |
| --- | --- | --- | --- | --- | --- | --- |
| (Intercept) | 6.798 | 0.363 | 28.806 | 18.712 | <0.001 |  |
| **Ax** | 3.398 | 0.532 | 31.019 | 6.391 | <0.001 | <0.001 |
| **Gy** | 0.658 | 0.520 | 30.095 | 1.266 | 0.215 | 0.513 |
| **Or** | 1.355 | 0.543 | 34.275 | 2.494 | 0.018 | 0.045 |
| **Pf** | 1.083 | 0.538 | 29.914 | 2.012 | 0.053 | 0.143 |

| Random Effect | Var. | SD |
| --- | --- | --- |
| Tank | 0.500 | 0.707 |
| Residual | 7.967 | 2.823 |

**Table 2.** Summary of model results for tail coloration (a* and b* values) measured at Week 3 after the start of Experiment II. Tadpoles were reared in the presence of no predators (the control group), water bug (*Appasus major*; Be), water scorpion (*Laccotrephes japonensis*; Ws), and fire belly newt (*Cynops pyrrhogaster*; Fn). Fixed effects include the corresponding estimates (Est.), standard errors (SE), degrees of freedom (df), t-values (*t*), and model P-values (*P*_m_). Post hoc comparisons were performed using Dunnett’s test to compare each predator treatment to the control group (*P*_D_). Variance (Var.) and standard deviation (SD) for the random effect (Tank) and residuals are also shown. The control group served as the reference level in all models.

Week 3
| Effect | Est. | SE | df | $t$ | $P_m$ | $P_D$ |
| --- | --- | --- | --- | --- | --- | --- |
| (Intercept) | -1.739 | 0.067 | 32.760 | -25.970 | <0.001 |  |
| **Be** | -0.031 | 0.094 | 31.019 | -0.336 | 0.739 | 0.954 |
| **Ws** | -0.097 | 0.092 | 29.311 | -1.053 | 0.301 | 0.578 |
| **Fn** | -0.086 | 0.092 | 29.619 | -0.933 | 0.358 | 0.655 |

| Random Effect | Var. | SD |
| --- | --- | --- |
| Tank | 0.015 | 0.123 |
| Residual | 0.258 | 0.508 |

$b^*$ value Week 3
| Effect | Est. | SE | df | $t$ | $P_m$ | $P_D$ |
| --- | --- | --- | --- | --- | --- | --- |
| (Intercept) | 5.871 | 0.216 | 28.429 | 27.198 | <0.001 |  |
| **Be** | 0.117 | 0.301 | 26.657 | 0.387 | 0.702 | 0.938 |
| **Ws** | 0.452 | 0.295 | 25.105 | 1.532 | 0.138 | 0.294 |
| **Fn** | 0.605 | 0.296 | 25.383 | 2.043 | 0.052 | 0.108 |

| Random Effect | Var. | SD |
| --- | --- | --- |
| Tank | 0.131 | 0.363 |
| Residual | 3.026 | 1.740 |

### Morphological Responses

In Experiment I, predators had minimal effects on body length (BL). The overall effect of predator treatment on BL was not statistically significant at either week (Week 2: *F* = 2.35, *P* = 0.075; Week 3: *F* = 2.35, *P* = 0.074), although a weak trend was observed. Tadpoles exposed to *P. flavescens* (Pf) developed slightly larger bodies than controls at the second week (Dunnett’s test, *P* = 0.048), but this difference was no longer significant at the third week (Table 3; Fig. 4). Similarly, although predator treatment had a significant overall effect on tail length, its effects were limited to certain groups or time points. When tail length was used as the response variable, models with body length and treatment as explanatory variables were selected as the final models at both the second and third weeks after the start of the experiment. In both models, tail length showed a strong positive association with body length (*P* < 0.001), indicating that larger tadpoles tended to have longer tails. In these models, predator treatment had a significant overall effect at both the second week (*F* = 6.76, *P* < 0.001) and the third week (*F* = 3.28, *P* = 0.023). Independent of BL, tadpoles in the Pf and *A. nigrofasciatus* (Ax) groups had shorter tails than controls at the second week (*P* < 0.001), but not at the third week. In contrast, tadpoles exposed to *Gynacantha japonica* (Gy group) showed reduced tail length only at the third week (*P* = 0.005).

**Fig. 4.**
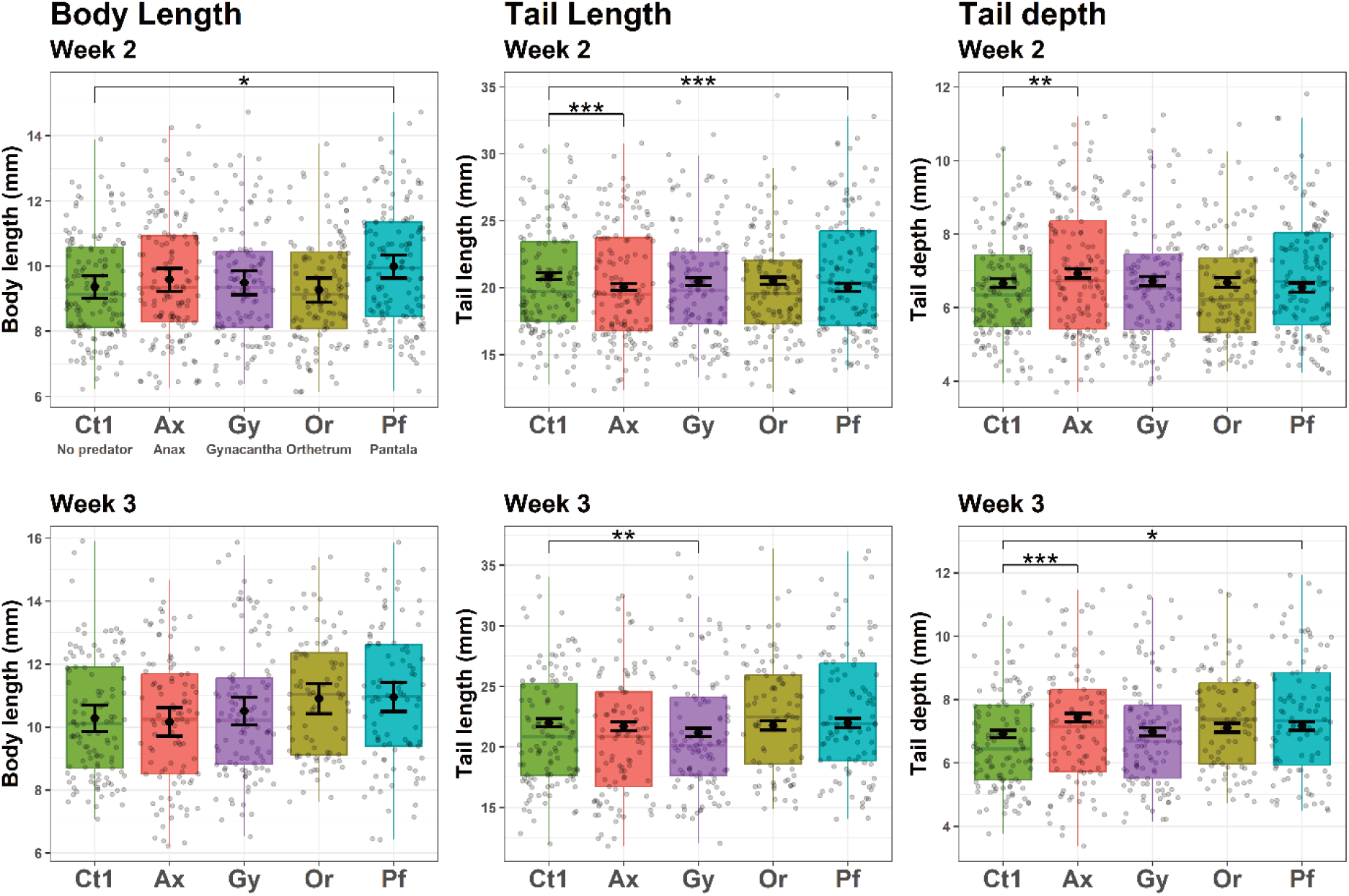
Box-and-whisker plots showing the distribution of tadpole morphological traits, BL (body length), TAL (tail length), and TD (tail depth) in Experiment I, using different dragonfly species. Tadpoles were reared in the presence of no predators (the control group; Ct1), *Anax nigrofasciatus* (Ax), *Gynacantha japonica* (Gy), *Orthetrum albistylum speciosum* (Or), or *Pantala flavescens* (Pf). Boxplots represent the raw data distributions, while black points and error bars indicate estimated marginal means and 95% confidence intervals derived from linear mixed-effects models (see Materials and Methods). The asterisks in the figure indicate that there were significant differences compared to the control group, which were led by Dunnett’s test. \**P* < 0.05, \*\**P* < 0.01, \*\*\**P* <0.001.

**Table 3.** Summary of model results for morphological traits (body length, tail length, and tail depth) measured at Week 2 and Week 3 after the start of Experiment I. Tadpoles were reared in the presence of no predators (the control group), *Anax nigrofasciatus* (Ax), *Gynacantha japonica* (Gy), *Orthetrum albistylum speciosum* (Or), or *Pantala flavescens* (Pf). Fixed effects include the corresponding estimates (Est.), standard errors (SE), degrees of freedom (df), t-values (*t*), and model P-values (*P*_m_). Post hoc comparisons were performed using Dunnett’s test to compare each predator treatment to the control group (*P*_D_). Variance (Var.) and standard deviation (SD) for the random effect (Tank) and residuals are also shown. The control group served as the reference level in all models.

| Effect | Est. | SE | df | *t* | *P<sub>m</sub>* | *P<sub>D</sub>* |
| --- | --- | --- | --- | --- | --- | --- |
| (Intercept) | 9.365 | 0.176 | 29.472 | 53.098 | <0.001 |  |
| **Ax** | 0.221 | 0.251 | 30.283 | 0.879 | 0.386 | 0.757 |
| **Gy** | 0.128 | 0.257 | 33.237 | 0.499 | 0.621 | 0.932 |
| **Or** | −0.098 | 0.257 | 33.152 | −0.381 | 0.706 | 0.964 |
| **Pf** | 0.626 | 0.253 | 31.049 | 2.472 | 0.019 | 0.048 |

| Random Effect | Var. | SD |
| --- | --- | --- |
| Tank | 0.077 | 0.277 |
| Residual | 3.047 | 1.746 |

**Tail length****Week 2**
| Effect | Est. | SE | df | *t* | *P<sub>m</sub>* | *P<sub>D</sub>* |
| --- | --- | --- | --- | --- | --- | --- |
| (Intercept) | −0.410 | 0.352 | 441.321 | −1.165 | 0.245 |  |
| Body length | 2.230 | 0.035 | 639.859 | 64.157 | <0.001 |  |
| **Ax** | −0.824 | 0.191 | 34.547 | −4.315 | <0.001 | <0.001 |
| **Gy** | −0.436 | 0.197 | 39.064 | −2.214 | 0.033 | 0.091 |
| **Or** | −0.376 | 0.197 | 38.784 | −1.910 | 0.063 | 0.178 |
| **Pf** | −0.860 | 0.194 | 36.388 | −4.439 | <0.001 | <0.001 |

| Random Effect | Var. | SD |
| --- | --- | --- |
| Tank | 0.006 | 0.079 |
| Residual | 2.416 | 1.554 |

**Tail depth****Week 2**
| Effect | Est. | SE | df | *t* | *P<sub>m</sub>* | *P<sub>D</sub>* |
| --- | --- | --- | --- | --- | --- | --- |
| (Intercept) | −1.229 | 0.141 | 373.721 | −8.733 | <0.001 |  |
| Body length | 0.827 | 0.014 | 642.972 | 61.130 | <0.001 |  |
| **Ax** | 0.266 | 0.088 | 31.452 | 3.029 | 0.005 | 0.009 |
| **Gy** | 0.061 | 0.090 | 34.327 | 0.683 | 0.499 | 0.860 |
| **Or** | 0.025 | 0.090 | 34.238 | 0.280 | 0.781 | 0.983 |
| **Pf** | −0.114 | 0.089 | 32.718 | −1.288 | 0.207 | 0.499 |

| Random Effect | Var. | SD |
| --- | --- | --- |
| Tank | 0.010 | 0.100 |
| Residual | 0.358 | 0.598 |

**Week 3**
| Effect | Est. | SE | df | *t* | *P<sub>m</sub>* | *P<sub>D</sub>* |
| --- | --- | --- | --- | --- | --- | --- |
| (Intercept) | 10.283 | 0.214 | 29.421 | 47.990 | <0.001 |  |
| **Ax** | −0.116 | 0.316 | 31.356 | −0.367 | 0.716 | 0.967 |
| **Gy** | 0.233 | 0.308 | 31.190 | 0.757 | 0.455 | 0.824 |
| **Or** | 0.627 | 0.326 | 36.584 | 1.921 | 0.063 | 0.174 |
| **Pf** | 0.673 | 0.319 | 30.756 | 2.111 | 0.043 | 0.116 |

| Random Effect | Var. | SD |
| --- | --- | --- |
| Tank | 0.082 | 0.288 |
| Residual | 4.089 | 2.022 |

**Week 3**
| Effect | Est. | SE | df | *t* | *P<sub>m</sub>* | *P<sub>D</sub>* |
| --- | --- | --- | --- | --- | --- | --- |
| (Intercept) | −2.121 | 0.409 | 316.445 | −5.182 | <0.001 |  |
| Body length | 2.287 | 0.036 | 488.929 | 63.557 | <0.001 |  |
| **Ax** | −0.307 | 0.257 | 29.473 | −1.192 | 0.243 | 0.560 |
| **Gy** | −0.810 | 0.251 | 29.271 | −3.224 | 0.003 | 0.005 |
| **Or** | −0.224 | 0.266 | 34.751 | −0.842 | 0.405 | 0.778 |
| **Pf** | −0.020 | 0.261 | 29.368 | −0.078 | 0.938 | 0.999 |

| Random Effect | Var. | SD |
| --- | --- | --- |
| Tank | 0.064 | 0.253 |
| Residual | 2.586 | 1.608 |

**Week 3**
| Effect | Est. | SE | df | *t* | *P<sub>m</sub>* | *P<sub>D</sub>* |
| --- | --- | --- | --- | --- | --- | --- |
| (Intercept) | −1.545 | 0.154 | 328.542 | −10.038 | <0.001 |  |
| Body length | 0.801 | 0.014 | 488.464 | 58.376 | <0.001 |  |
| **Ax** | 0.519 | 0.091 | 27.792 | 5.723 | <0.001 | <0.001 |
| **Gy** | 0.068 | 0.089 | 28.102 | 0.763 | 0.452 | 0.821 |
| **Or** | 0.200 | 0.095 | 34.000 | 2.119 | 0.042 | 0.113 |
| **Pf** | 0.253 | 0.092 | 28.144 | 2.751 | 0.010 | 0.022 |

| Random Effect | Var. | SD |
| --- | --- | --- |
| Tank | 0.004 | 0.061 |
| Residual | 0.380 | 0.617 |

The impact of predator treatment on tail depth was more pronounced. When tail depth was used as the response variable, BL and treatment were chosen as explanatory variables at both the second and third weeks after the start of the experiment. In both models, tail depth also showed a strong positive association with BL (*P* < 0.001), and the effects of predator treatments were interpreted independently of body size. Predator treatment had a strong overall effect on tail depth at both the second week (*F* = 4.86, *P* = 0.003) and the third week (*F* = 9.60, *P* < 0.001). Post-hoc tests revealed that tadpoles in the Ax group developed deeper tails than controls at both time points (*P* < 0.01). Moreover, tadpoles in the Pf group also developed deeper tail fins than controls at the third week (*P* = 0.022). A similar trend was observed in the *O. albistylum* (Or) group, although this difference did not reach statistical significance (*P* = 0.113).

In Experiment II, predators had minimal effects not only on BL, but also on tail morphology. Predator treatment had no significant effect on BL (Week 2: *F* = 1.82, *P* = 0.168; Week 3: *F* = 2.18, *P* = 0.118). For tail length, models including both BL and treatment as explanatory variables were chosen at the second week. In this model, tail length showed a strong positive association with BL (*P* < 0.001), and predator treatment had a significant overall effect (*F* = 7.43, *P* < 0.001). Post-hoc tests revealed that a significant reduction in tail length was observed in the Ferocious water bug (Be) group (*P* < 0.001; Fig. 5). At the third week, however, models with only BL were selected, indicating that the effect of the treatment on tail length appeared limited. Similarly, for tail depth, models with only BL were selected in both weeks, and tail depth was positively associated with body length (*P* < 0.001). Post-hoc tests also confirmed that tail depth was largely unaffected, with no significant differences observed. Thus, tadpoles in all non-odonate predator treatments were not induced to develop significantly deeper tail fins compared to the control group.

**Fig. 5.**
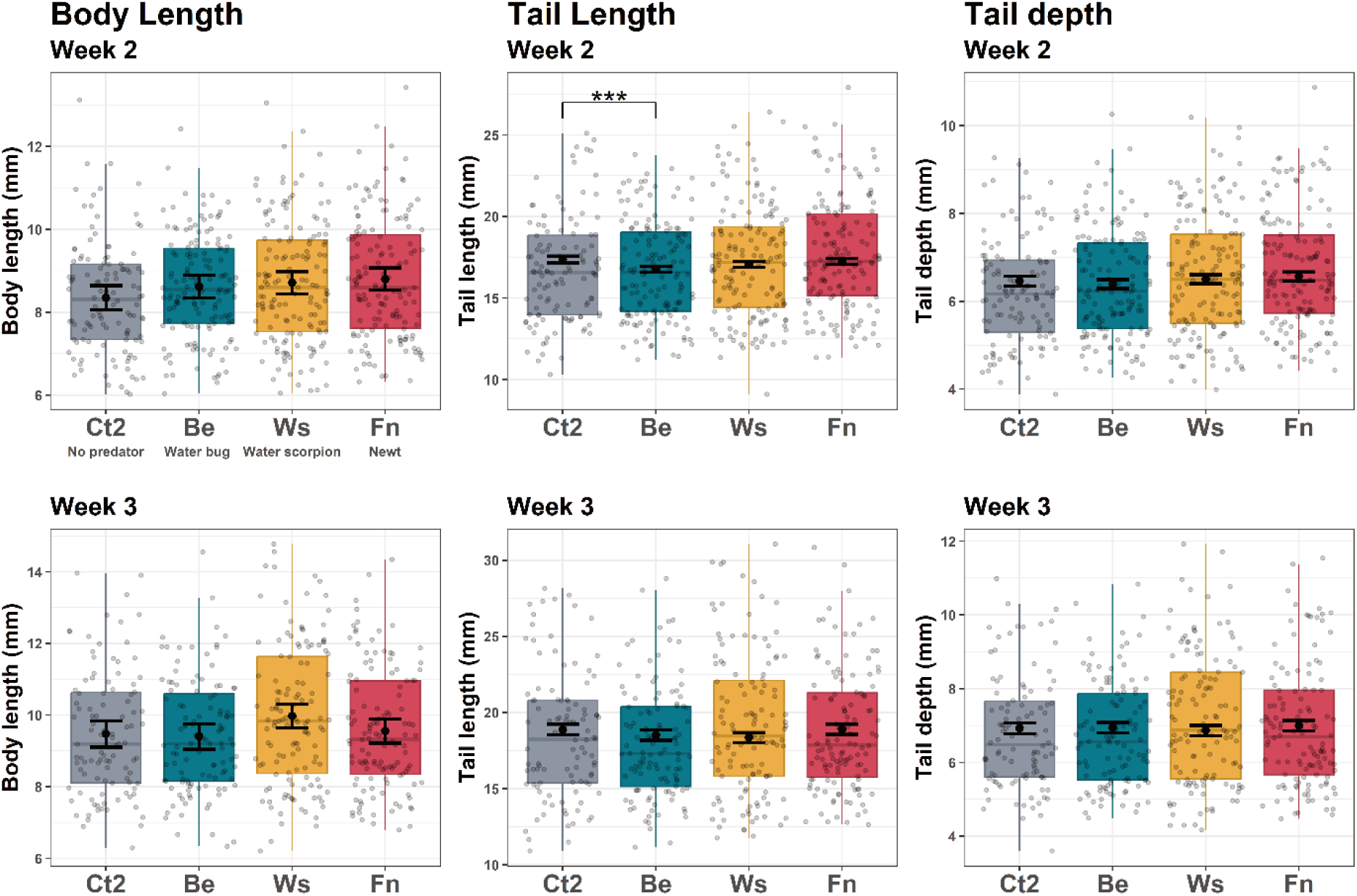
Box-and-whisker plots showing the distribution of tadpole morphological traits, BL (body length), TAL (tail length), and TD (tail depth) in Experiment II, using non-odonate predators. Tadpoles were reared in the presence of no predators (the control group; Ct2), water bug (*Appasus major*; Be), water scorpion (*Laccotrephes japonensis*; Ws), and fire belly newt (*Cynops pyrrhogaster*; Fn). Boxplots represent the raw data distributions, while black points and error bars indicate estimated marginal means and 95% confidence intervals derived from linear mixed-effects models (see Materials and Methods). The asterisk in the figure indicates a significant difference compared to the control group based on Dunnett’s test. \*\*\**P* <0.001.

The orange coloration and deeper tail morphology tended to be co-expressed within individuals. Significant, though weak, positive correlations between b* values and tail depth were detected in all three odonate predator treatments that induced changes in either trait by the third week (Pf: *r* = 0.393, *P* < 0.001; Ax: *r* = 0.294, *P* = 0.003; Or: *r* = 0.277, *P* = 0.011).

## DISCUSSION

### Tail Coloration in Asian Species of Dryophytes

Our experiments revealed that tadpoles of *Dryophytes leopardus* developed induced changes in tail coloration as well as tail shape in response to dragonfly nymphs. Although tail coloration has rarely been documented in this species or in its close relative, *Dryophytes japonicus*, its occurrence has been reported from several locations in Japan, including Hokkaido, Shiga, Hyogo, Hiroshima, Shimane, Kouchi and Ehime prefectures (Doi et al., 2005), as well as from our collection site in Kyoto (Noda, personal observation). The phenotypes induced by *Anax nigrofasciatus*, i.e., deep tail fins and bright orange tails, closely resemble those previously observed in other species in *Dryophytes* in the New World, Cope’s Gray Treefrog (*Dryophytes chrysoscelis*; McCollum and Van Buskirk, 1996), Pine Woods Treefrog (*Dryophytes femoralis*; LaFiandra and Babbitt, 2004), and Gray Treefrog (*Dryophytes versicolor*; Van Buskirk and McCollum, 1999). Thus, our findings represent the first documented case of a predator-induced tail coloration phenotype in Asian species of *Dryophytes*.

Our discoveries of the plasticity in the tail coloration in the Asian species of *Dryophytes* suggests that the trait is symplesiomorphically shared between the North American and ancestral Asian lineages, and has been inherited by present-day Asian species, including *D. leopardus*. An ancestral stock of *Dryophytes* is inferred to have migrated from North America to East Asia across the Bering Land Bridge during the Miocene (Duellman et al. 2016; Dufresnes et al., 2016). Present-day Asian species of *Dryophytes* are monophyletic, having diversified from this ancestral stock. It is plausible that tail coloration has persisted as a phenotypic plasticity in both North American and East Asian species of *Dryophytes* due to the consistent importance of dragonfly nymphs as predators.

It remains unknown whether tail coloration can be induced in the three other Asian species, *Dryophytes flaviventris*, *Dryophytes immaculates*, and *Dryophytes suweonensis* (see Borzée et al., 2020). In North America, Pine Barrens Treefrog, *Dryophytes andersonii* are induced to have a darker tail, not an orange tail, by *Anax junius*, which induces orange tails in other species of *Dryophytes* (Kruger and Morin, 2020). Furthermore, Squirrel Treefrog, *Dryophytes squirellus* exhibits no morphological changes in response to odonate species (McCoy and Bolker 2008; Baker, 2021). These cases indicate that the plastic responses in *Dryophytes* have diversified, and some may have lost the ability to develop orange tails. Future studies are needed to elucidate the interspecific variation in plastic responses to predation risk and the evolutionary causes underlying this variation.

### Differences in the Extent of Coloration by Predators

Our experiments demonstrated that the extent and timing of induced responses varied depending on the dragonfly species reared with tadpoles. Two weeks after the start of the experiment, nymphs of *A. nigrofasciatus* induced both deep fin and tail coloration in tadpoles. Additionally, by the third week, similar but weaker responses were observed in the other predator treatments: *Pantala flavescens* also induced deep tail and *Orthetrum albistylum speciosum* induced tail coloration. Furthermore, although weak, some tadpole individuals exposed to *P. flavescens* showed tail coloration, and those exposed to *O. albistylum speciosum* also tended to have deeper tails. These results indicate that the defense traits (coloration and deep tail) were induced by the nymphs of *A. nigrofasciatus* more rapidly and strongly, but similar responses were also elicited by the other two predator species. Furthermore, the significant positive correlation observed between the b* value and tail depth suggests that these traits tended to be co-expressed in a coordinated manner. These findings support the idea that the expression of these anti-predator defenses is governed by a shared, integrated strategy, whose intensity varies depending on predator identity.

The differences in the degree and rate of responses induced by different dragonfly species may reflect varying levels of predation risk for tadpoles posed by each species. Plastic phenotypes with significant tail coloration should result in trade-offs because of energetic costs associated with synthesizing pigmentation (Grether et al., 2001) or costs due to the increased vulnerability to mismatched predators. Consequently, tadpoles are expected to minimize these costs by developing effective defense traits for each predator as much as possible based on the perceived predation risk (Touchon and Warkentin, 2008; Innes-Gold et al., 2019). The predation risk for tadpoles would be different among dragonfly species according to (1) the foraging style of dragonfly nymphs, (2) their color vision, and (3) the total amount of intake related to their body size and occurrence period.

Firstly, the four dragonfly species used in our experiment exhibit different foraging styles. According to the classification by Corbet (1999), *A. nigrofasciatus* and *G. japonica* are climbers, sit-and-wait predators on aquatic vegetation, while *P. flavescens* is a sprawler, foraging while moving along the bottom, and *O. albistylum speciosum* is a burrower, residing within the substrate and ambushing prey. For sit-and-wait predators, there is often a time lag between identifying prey and launching an attack, during which they can choose which part of tadpole’s body to target. Therefore, the lure effect of tadpole tails is likely to attract ambush predators more effectively than those that actively chase their prey. In addition, as *A. nigrofasciatus* prefers habitats abundant in floating-leaved plants (Ozono et al., 2021), it can move freely along the plant structures from roots anchored at the bottom to leaves extending above the water surface. As a result, the risk of predation on tadpoles could be present throughout the entire water column, from the bottom to the surface, making behavioral responses like moving to a specific depth insufficient as a means of avoiding predation.

Conversely, *P. flavescens* and *O. albistylum speciosum* primarily forage at the bottom, so tadpoles may evade these predators by relocating to higher water layers. However, such avoidance behavior often leads to a decrease in growth rate due to reduced foraging efficiency (Teplitsky et al., 2003); i.e., for tadpoles that primarily feed on detritus, avoiding the bottom layer may result in malnutrition. The combination of moderate morphological responses with behavioral responses (the “double defense” strategy; Dewitt and Hucko, 1999) may help mitigate the impact of malnutrition from behavioral changes while minimizing the costs of defensive traits.

Secondly, differences in eyesight, particularly color vision among dragonfly species, may influence the extent of the tail coloration changes in the tadpoles. Pritchard (1966) noted a correlation between foraging habit and compound eye size in dragonfly nymphs, with climbers typically possessing larger compound eyes compared to sprawlers and burrowers. He also stated that the climbers, such as species in *Anax*, rely heavily on vision for prey detection, whereas other foraging types depend more on tactile stimuli. Hence, it is plausible that the lure effect of the brightly colored tails is more effective for climbers, which are more visually oriented. Furthermore, Futahashi et al. (2015) investigated the genetic basis of color vision in dragonflies and redocumented the number of opsin genes and the characteristic spatiotemporal expression patterns in the nymphs of various species, including those of the burrower *O. albistylim* and the climber *A. parthenope*, a species closely related to *A. nigrofasciatus* used in our experiment. They found that dragonflies possess a greater number of opsin genes than other insects, and *A. parthenope* possesses and expresses a greater number of opsin genes, particularly those sensitive to long-wavelength light. Although the functions of these diverse genes are not fully understood, this suggest that species of *Anax* may have enhanced abilities to perceive reddish colors, potentially enabling tadpoles with bright orange tails to more effectively avoid predation by using their tails as lures. Thus, the larval antipredator defense of *D. leopardus*, which involves tail coloration, may have been driven by the sensory systems of predatory dragonfly nymphs, and subsequently evolved specifically against color-sensitive predators, represented by dragonfly nymphs of *Anax*.

Thirdly, the variation in the degree of induced responses may reflect the relative predation risks associated with body sizes or occurrence periods of respective odonate predators. In nature, it is generally difficult to determine which predators exert higher predation pressure on tadpoles, and few studies have investigated such differences. However, during the rearing process, we observed that the nymphs of *A. nigrofasciatus* exhibited higher food demands and displayed more frequent attacks on tadpoles compared to other species.

Additionally, they are approximately twice the size of the nymphs of *P. flavescens* and *O. albistylum speciosum* (see Ishida et al., 1988), suggesting that size-related predation pressures could influence the necessity and effectiveness of tadpole morphological responses. However, tadpoles did not exhibit significant responses to *G. japonica*, which is similar in size to *A*. *nigrofasciatus* (see Ishida et al., 1988). This indicates that tadpoles likely respond not only to size-related predation pressure but also to other characteristics of dragonfly species, such as foraging style or habitat overlap, thereby modulating the extent of induced responses. Since the habitat of nymphal *G. japonica* overlaps with that of tadpoles of *D. leopardus* (Nakagawa et al., 2004), the lack of an apparent induced response may be due to the limited period in which they co-occur. This species grows more rapidly compared to other species in Aeshnidae in Japan, occurring in aquatic environments only from May to August (Ozono et al., 2021).

Considering that the nearly-metamorphosing nymphs cease foraging, the period during which this species poses a predation threat to *D. leopardus* is even shorter. This brief coexistence period suggests that this dragonfly species may not be an important predator for the tadpoles.

In addition to the tail coloration and depth we focused on, a tendency toward larger body lengths and shorter tail lengths compared to those in the control group was observed, although not statistically significant in most treatment groups. These changes may represent general responses to predator treatments, as noted in previous studies. The increase in body length could be attributed to enhanced availability of food resources resulting from leftovers and feces of caged predators (e.g., McIntyre et al., 2004; Richardson, 2006). Similarly, shorter tail lengths in the presence of predators have been reported in several frog species (Laurila et al., 2002; LaFiandra and Babbitt, 2004).

### Responses Toward Non-Odonate Predators

Tadpoles of *D. leopardus* showed neither conspicuous tail coloration nor deep fin when exposed to newts or aquatic insects other than dragonfly nymphs in our experiment, likely because the costs of expressing these defenses outweighed their benefits for avoiding predation. Although Experiment II did not include a positive control (i.e., treatment with dragonfly nymphs, such as *A*. *nigrofasciatus*), we believe that the absence of color induction was unlikely caused by experimental failure, because preliminary experiments demonstrated the high reproducibility of the induced responses when using dragonfly nymphs. Therefore, our results suggest that changes in body color and tail depth are not strongly induced by these predators and may not serve as effective predation avoidance strategies against them. An important point in understanding why tadpoles did not show tail coloration is that defensive traits come with physiological costs and may increase predation pressure from non-focal predators. If changes in tail color and shape do not significantly contribute to non-odonate predator avoidance, tadpole should minimize these disadvantages by refraining from developing these antipredator defenses.

Given that the non-odonate predators do not target specific body parts of their prey during attacks, the strategy of using colored deep tails as lures may be ineffective and provide only limited benefits against them. For instance, diving beetles, water bugs, and water scorpions capture prey by grasping the entire body with their arms, so the tails of tadpoles should not serve as lures for them. Additionally, they use tactile, visual, and olfactory senses for locating prey (Babbitt and Jordan, 1996; Kehr and Gómez, 2009; Mogali et al., 2019; Nowińska and Brożek, 2019), and possess well-developed organs for detecting auditory and mechanical signals (Yager, 1999; Cocroft and Rodríguez, 2005). Previous studies have reported predator-induced changes in larval morphology in response to water bugs (Benard, 2006; McCoy, 2007; Michel, 2012), but few have reported effects on larval tail coloration (McIntyre et al., 2004; Touchon and Warkentin, 2011). Similarly, newts and salamanders, both as larvae and adults, are active foragers that swallow their prey whole like fish (Wilson et al., 2005; Kishida and Nishimura, 2007). Consequently, the tails of the tadpoles are unlikely to function as a lure for these predators. Newts tend to rely on olfactory cues as well as vision for food recognition (Copeland, 1913; Lindquist and Bachmann, 1982; Schwarz et al., 2020). Few studies have reported that tadpoles exhibit induced defensive traits in response to these predators, and none have documented induced changes in body coloration (Kishida and Nishimura, 2004; Hettyey et al., 2011; Urban et al., 2017). These facts support the hypothesis that the brightly colored tails of *D. leopardus* do not function as lures against the non-odonate predators, and that, as a result, tadpoles do not express costly defensive traits in response to them.

The lack of induction of defensive phenotypes may also be attributed to physiological costs associated with developing tail coloration, although the magnitude of such costs in tadpoles remains poorly understood. Since most animals are unable to biosynthesize carotenoids, the pigments thought to be causing the bright orange coloration, they metabolize and accumulate these pigments by absorbing them from their diet (Bertrand et al., 2006; Thibaudeau and Altig, 2012). Thus, the allocation costs of expressing coloration by transporting these pigments may not be as high as commonly assumed. On the other hand, the potential for increased vulnerability to non-focal predators may also interfere with the expression of defensive traits (Innes-Gold et al., 2019). This disadvantage may sometimes outweigh the energetic costs of expressing a defensive phenotype. Carfagno et al. (2011) found that tadpoles of *Acris* exhibit darker a tail spot, which is considered a defensive trait against dragonfly nymphs, even in the absence of predators. However, the presence of fish predators induces the loss of this spot. This suggests that the cost of expressing defensive traits may be lower than typically expected, and the disadvantage of increased vulnerability to mismatched predators may prevent these phenotypes from being constitutively expressed.

Similarly, in tadpoles of *D. leopardus*, the elevated risk of predation from non-odonate predators, considered non-focal predators, due to expressing bright tail coloration, rather than its energetic cost, could explain why they did not exhibit deep and brightly-colored tail in response to these predators. These defense traits against dragonfly nymphs may not only be ineffective in preventing predation by non-odonate predators but could also increase the potential vulnerability to them. For example, the deepened tail may increase the vibrations produced during swimming, thereby heightening predation pressure on tadpoles by tactile or olfactory predators (i.e., water bugs, water scorpions, and newts). Additionally, for other predators such as fish, tail coloration may make the tadpoles more vulnerable by increasing their visibility. However, further experiments are warranted to assess and compare the effectiveness of these defense traits toward each predator, including evaluating how tadpole coloration influence their survival rate in the presence of various aquatic predators.

## ACKNOWLEDGMENTS

We followed the ASIH “Guidelines for Live Amphibians and Reptiles in Field and Laboratory Research” (https://bit.ly/ASIH_Herps). The described methods were conducted in compliance with the Regulation on Animal Experimentation at Kyoto University. Because the aim of the experiments was to induce the anti-predator phenotypes of tadpoles, we needed to use live tadpoles and some of them were eaten by their predators in the experiments.

However, the process was the same as a predation event occurring in nature, which did not cause any unnatural pain to the tadpoles. We thank the Experimental Farm, Graduate School of Agriculture, Kyoto University, for providing the research environment, and Mr. Kagata, Mr. Yasui, and Mr. Sakakibara for their assistance with observations. This work was supported by JST SPRING, Grant Number JPMJSP2110.

## Notes

### Competing Interest Statement

The authors have declared no competing interest.

### Summary of Updates

This version updates the manuscript to the accepted version for publication in Ichthyology & Herpetology. The following changes have been made: (1) the text has been revised for clarity and grammar; (2) figures and tables have been updated to the final versions; (3) minor corrections to data presentation and references have been incorporated. A statement has been added to indicate that the article has been accepted for publication in Ichthyology & Herpetology.

